# Ecological context structures duplication and mobilization of antibiotic and metal resistance genes in bacteria

**DOI:** 10.64898/2026.02.03.703612

**Authors:** Eric Tran, Priscilla N. Xu, Raquel Assis

## Abstract

Antibiotic resistance is a global challenge driven by the persistence and spread of resistance genes across ecological contexts. While mobile genetic elements (MGEs) facilitate horizontal gene transfer, gene duplication represents an additional mechanism through which resistance genes can be amplified, diversified, and maintained under selection. How these processes interact across environments remains poorly understood. Here, we examined genome-level patterns of resistance gene abundance, duplication, and mobilization across clinical, agricultural, and wastewater settings, focusing on both antibiotic resistance genes (ARGs) and metal resistance genes (MRGs). Resistance gene profiles were strongly structured by environment, with distinct duplication patterns emerging across sources. Duplicate genes were frequently associated with MGEs, although the strength of this relationship varied by resistance type and ecological context. Despite frequent co-occurrence of ARGs and MRGs, their duplication and mobilization dynamics were not uniformly coupled at the genome level. Together, these findings highlight gene duplication as a context-dependent contributor to resistance evolution and underscore the importance of ecological setting in shaping how resistance genes persist and spread across microbial communities.

## Introduction

Antibiotic resistance poses a growing global threat to public health, driving increases in infection-related hospitalizations, fatalities, and healthcare costs as a consequence of decades of antibiotic overuse and overprescription (Llor and Bjerrum, 2014). The World Health Organization projects that antibiotic-resistant infections could cause up to 10 million deaths annually by 2050 (World Health Organization, 2019). Antibiotic resistance genes (ARGs) are widespread in both clinical and environmental settings, with hotspots including hospitals, livestock farms, and wastewater treatment facilities (Berendonk et al., 2015; Larsson and Flach, 2022). Understanding the mechanisms that spread and amplify these genes across environments is critical for developing effective intervention strategies.

Many ARGs reside on mobile genetic elements (MGEs), such as plasmids and transposons, which are key drivers of horizontal gene transfer (HGT) (Rocha et al., 2022; Tokuda and Shintani, 2024). These elements enable resistance genes to move across bacterial strains, species, and ecological boundaries, accelerating the dissemination of resistance within and between microbial communities (Partridge et al., 2018). By allowing bacteria to acquire pre-existing resistance genes rather than waiting for de novo mutations, MGEs substantially increase the rate at which populations can adapt to antibiotic pressure (Ochman et al., 2000; Kumavath et al., 2025). They also help maintain ARGs in microbial communities even when selection is weak, through processes such as broad host-range transfer and stabilizing genomic architectures that favor persistence (San Millan and MacLean, 2017; Lopatkin et al., 2017). As a result, MGEs create shared reservoirs of resistance across clinical, agricultural, and environmental settings and contribute to the long-term evolutionary flexibility of microbial genomes.

Because antibiotic-rich environments impose strong and recurrent selective pressures on bacterial populations, their evolution depends on mechanisms that generate new genetic variation. Gene duplication is a major source of such variation, producing additional gene copies that can increase dosage, modify regulatory potential, or furnish opportunities for functional innovation during adaptation to environmental stressors (Ohno, 1970; Andersson and Hughes, 2009; Sandegren and Andersson, 2009; Bratlie et al., 2010; Kondrashov, 2012). In the context of resistance evolution, duplicated ARGs can elevate resistance levels or create substrate-specific variants, making duplication directly relevant to adaptation in antibiotic-rich settings. Importantly, MGEs can mediate duplication events under selective pressure, including the duplication of ARGs in antibiotic-rich settings (Maddamsetti et al., 2024). Yet despite these insights, we still lack a broader understanding of how duplication operates across different classes of resistance genes and ecological conditions.

Metal resistance genes (MRGs) are particularly informative for addressing this gap because they are widespread in natural and human-impacted environments, and the metals they confer resistance to impose strong selective pressures on microbial populations (Bruins et al., 2000; Pal et al., 2022). Moreover, MRGs are often genetically linked to ARGs, indicating that metals and antibiotics may shape resistance evolution through coupled processes (Baker-Austin et al., 2006; Li et al., 2017; Gillieatt and Coleman, 2024). Their shared mobilization by MGEs makes MRGs an ideal system for studying how HGT and gene duplication jointly shape microbial adaptation. This linkage also provides a framework for comparing how duplication dynamics differ between antibiotic and metal resistance systems.

Historically, the study of MGEs has been constrained by the limitations of short-read whole-genome sequencing (Kingsford et al., 2010). Although short-read sequencing provides valuable information about genomic content, it is often insufficient to resolve the large, repetitive, and structurally complex regions typical of MGEs (Partridge et al., 2018). Consequently, the genomic architecture of MGEs and their associated genes is often fragmented or misassembled, obscuring patterns of gene duplication and mobilization (Arredondo-Alonso et al., 2017). Advances in long-read sequencing now allow the generation of complete, contiguous genomes, providing an unprecedented opportunity to characterize MGEs and their evolutionary dynamics with greater accuracy and resolution (Zhao et al., 2023).

In the present study, we leverage complete bacterial genomes generated using long-read sequencing to investigate how MGEs shape patterns of antibiotic and metal resistance across four ecological contexts representing clinical, agricultural, and urban wastewater environments. By analyzing complete genomes from these heterogeneous selective landscapes, we demonstrate that ARGs are more prevalent in environments with high antibiotic burden and that duplicate genes are mobilized more frequently than single-copy genes, underscoring their role in HGT. We also find that patterns of antibiotic and metal resistance vary systematically among ecological settings, revealing how environmental pressures shape the evolutionary dynamics of microbial genomes.

## Results

### Environmental variation in resistance gene abundance and duplication

We assessed how ecological context shapes resistance evolution by comparing the abundance of ARGs and MRGs across four distinct environments: Barnes-Jewish Hospital, Cambridge University Hospitals, livestock farms, and wastewater treatment works (WwTW; see *Methods*). ARG abundance varied markedly among these environments (Fig. 1A). Clinical isolates from Barnes-Jewish Hospital and Cambridge University Hospitals harbored the highest ARG burdens, livestock isolates exhibited moderately elevated ARG counts, and WwTW isolates showed the lowest ARG abundance, despite representing a convergence point for multiple sources of antibiotic input (Economou and Gousia, 2015; Manyi-Loh et al., 2018; Akhter et al., 2024). In contrast, MRG abundance displayed a distinct environmental pattern (Fig. 1B), with the highest levels observed in WwTW isolates and broadly similar abundances across the remaining environments. These patterns were generally robust to species composition: ARG abundance trends were preserved when *Escherichia coli* was excluded or analyzed alone (Figures S1A and S2A), although limited sample sizes for non–*E. coli* taxa reduced power to detect MRG abundance differences in WwTW isolates.

**Figure 1.**
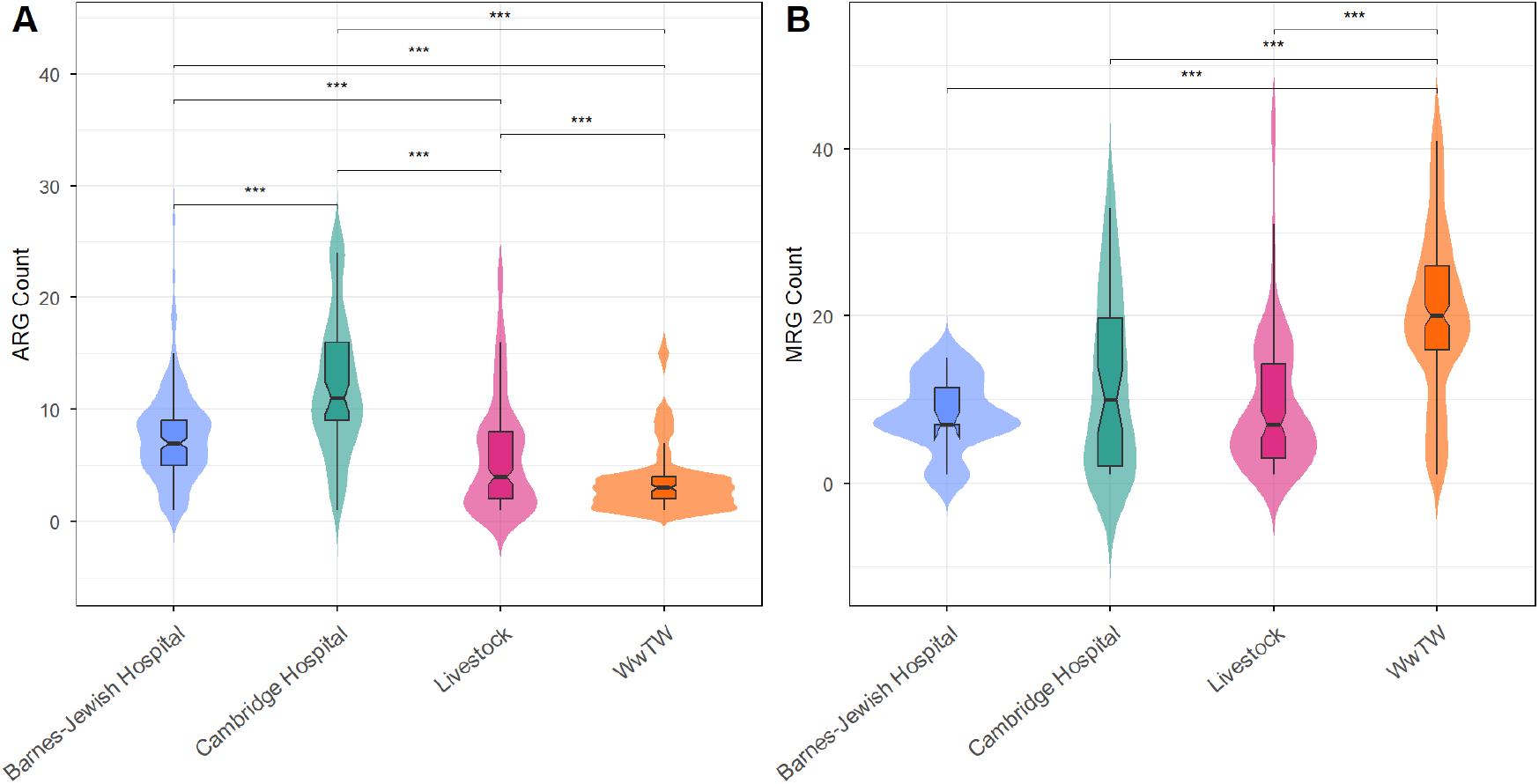
Abundance of ARGs (**A**) and MRGs (**B**) per genome across the four environments. Violin plots show distributions of gene counts, with embedded boxplots indicating the median and interquartile range. \**p* < 0.05, \*\**p* < 0.01, \*\*\**p* < 0.001 (see *Methods*).

To assess whether environmental context also shapes resistance gene duplication, we quantified the frequencies of duplicated ARGs and MRGs across environments (Fig. 2). Patterns of ARG duplication were significantly different across environments (*p* = 1.35 × 10^−7^, Fisher’s exact test; see *Methods*) and closely mirrored overall ARG abundance, with the highest frequencies observed in isolates from the two hospital settings and lower frequencies in livestock and WwTW isolates. Similarly, duplication patterns for MRGs reinforced the strong environmental structure observed for MRG abundance (*p* = 3.32 × 10^−34^, Fisher’s exact test; see *Methods*). Duplicate MRGs were most prevalent in WwTW isolates, followed closely by Cambridge Hospital isolates, whereas Barnes-Jewish Hospital and livestock isolates exhibited substantially lower frequencies. Collectively, these results demonstrate that ARGs and MRGs exhibit contrasting patterns of abundance and duplication across environments.

**Figure 2.**
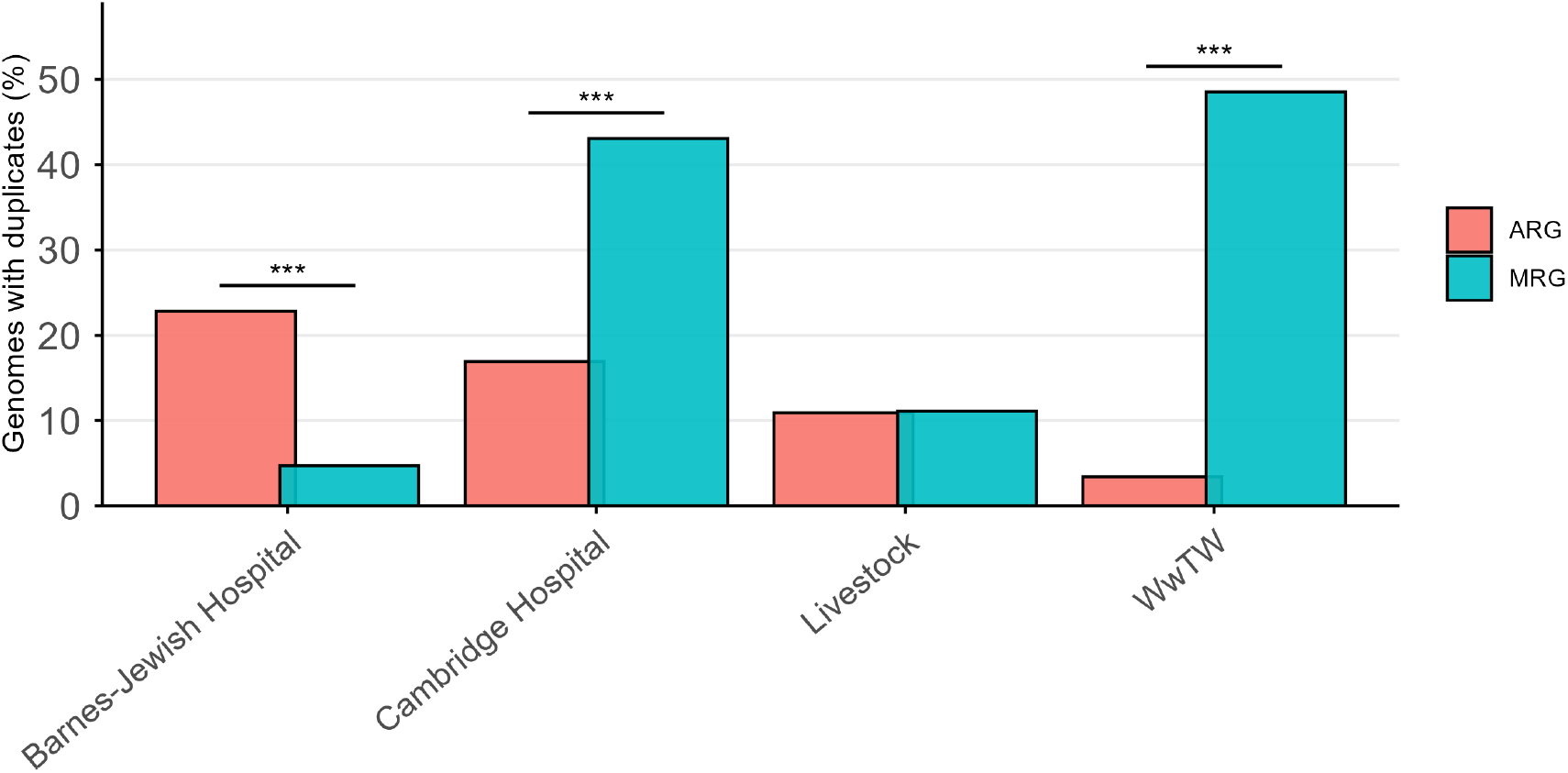
Percentage of genomes containing at least one duplicate resistance gene in each of the four environments. ARG and MRG proportions were calculated independently, so a genome may contribute to both categories. \**p <* 0.05, \*\**p <* 0.01, \*\*\**p <* 0.001 (see *Methods*).

### Gene duplication and mobilization of resistance genes across MGEs

To explore whether gene duplication facilitates the spread of resistance, we examined the association of single-copy and duplicate ARGs and MRGs with MGEs. Duplicate genes were disproportionately associated with MGEs, showing significant enrichment in 84% of the genomes analyzed (769 out of 913; Table S1). This enrichment is consistent with previous observations that duplicate genes are frequently carried by MGEs (Maddamsetti et al., 2024) and suggests that duplicate genes are preferentially mobilized relative to single-copy genes.

We next analyzed the distribution of single-copy and duplicate ARGs and MRGs across MGE classes (Fig. 3). Most ARGs and MRGs were associated with an MGE, with plasmids accounting for the majority of mobilized genes across all environments. While insertion sequences (ISs), composite transposons, and unit transposons comprised most of the remaining mobilized genes, phage-associated genes represented only a small fraction of the total pool. Interestingly, miniature inverted-repeat transposable elements (MITEs) were observed exclusively among single-copy genes from Cambridge Hospital and livestock isolates, and no ARGs or MRGs were detected on integrative conjugative elements (ICEs).

**Figure 3.**
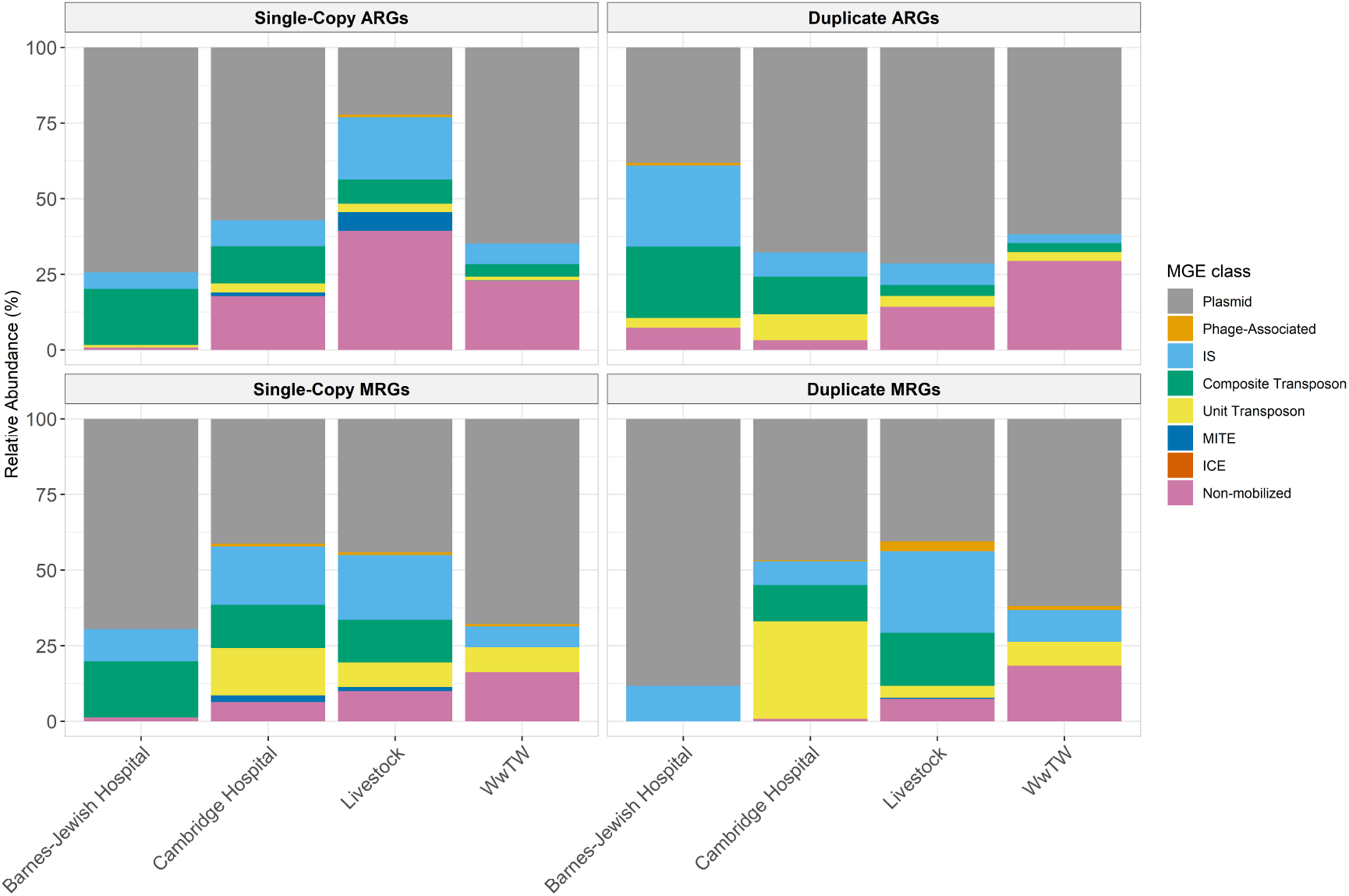
Relative abundance of MGEs associated with ARGs (top) and MRGs (bottom) across four environments. Left panels show single-copy genes, and right panels show duplicate genes. Bars are stacked by MGE type, with values normalized to the percentage of genes per genome.

Although duplicate genes were generally more frequently associated with MGEs than single-copy genes (Fig. 3, left vs. right panels), this enrichment varied across environments when considering ARGs and MRGs specifically. Duplicate ARGs showed higher mobilization in Cambridge Hospital and livestock isolates, whereas Barnes-Jewish Hospital and WwTW isolates exhibited similar or higher mobilization of single-copy ARGs. A comparable pattern was observed for MRGs in WwTW, although differences were modest. Overall, these results indicate that duplicate resistance genes are typically mobilized via plasmids, with extent and pattern of mobilization varying across environments and gene classes.

### Source-Specific Profiles of Duplicate ARG and MRG Classes

If gene duplication is driven by positive selection for environmental adaptation, the composition of duplicate ARGs and MRGs should reflect the selective pressures characteristic of each environment. To address this hypothesis, we quantified the total number of ARGs and MRGs from each source, grouped them by resistance class, and calculated the difference between the proportions of duplicate and single-copy genes within each class (Δ_*D*−*S*_), with positive values indicating enrichment of duplicate genes (see *Methods*).

We observed substantial environmental variation in the distribution of duplicate ARGs and MRGs across resistance classes (Fig. 4, Table S2). Among ARGs, Barnes-Jewish Hospital isolates were significantly enriched only for *β*-lactam resistance duplicates (Δ_*D*−*S*_ = 26.72%), consistent with their extended-spectrum *β*-lactamase (ESBL)-producing phenotype. In contrast, Cambridge Hospital isolates showed significant enrichment exclusively in aminoglycoside-quinolone duplicate genes (Δ_*D*−*S*_ = 21.66%), despite positive Δ_*D*−*S*_ values observed in additional classes. Outside clinical settings, livestock isolates exhibited significant enrichment in aminoglycoside ARGs (Δ_*D*−*S*_ = 37.82%), whereas WwTW isolates did not show statistically significant enrichment in any resistance class, despite several classes displaying positive Δ_*D*−*S*_ values.

**Figure 4.**
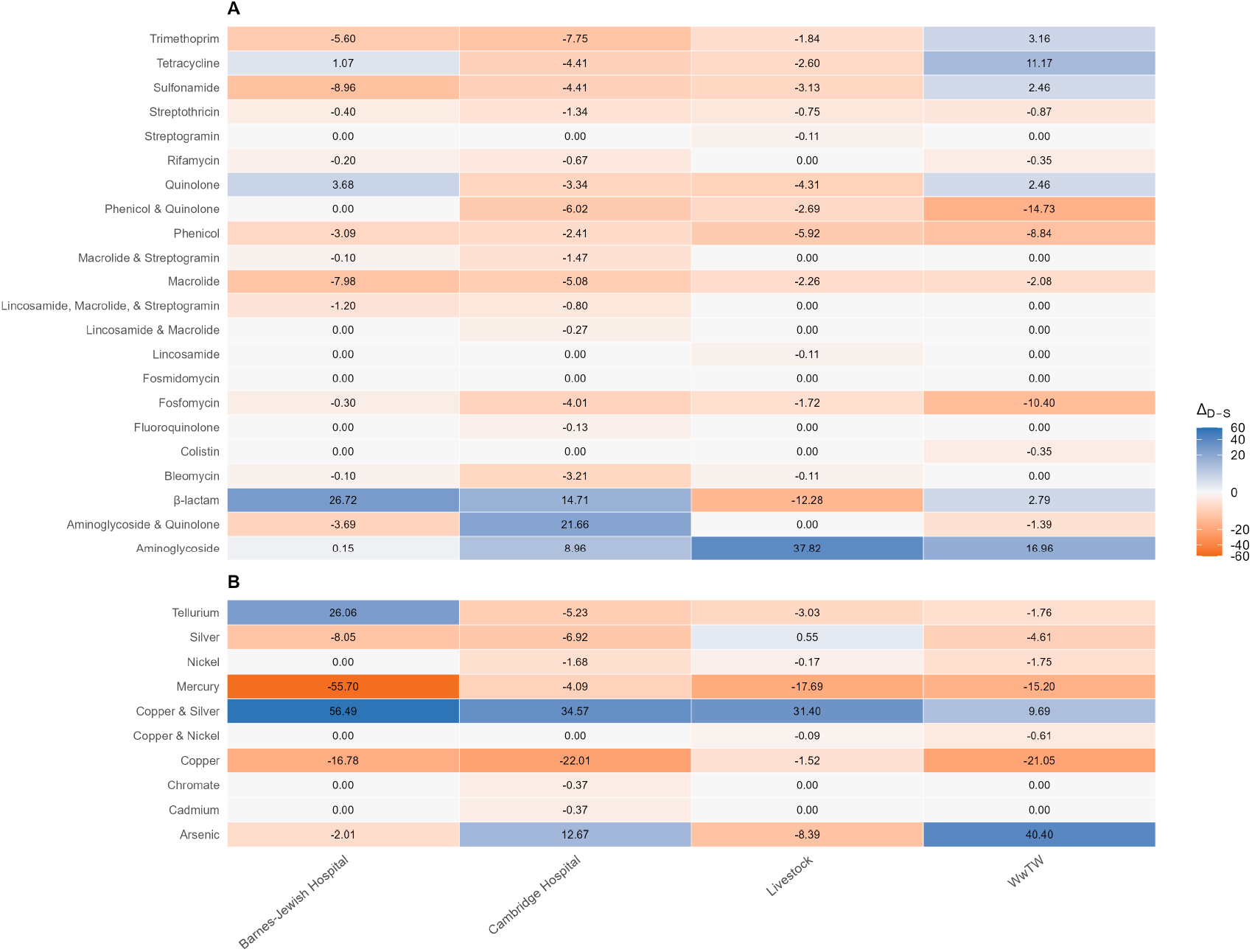
Heatmaps showing differences in the frequencies of duplicate and single-copy genes (Δ_*D*−*S*_) for ARG (**A**) and MRG (**B**) classes across the four environments. Positive values (blue) indicate enrichment of duplicate genes, while negative values (orange) indicate enrichment of single-copy genes. Statistical test results are provided in Table S2.

For MRGs, copper–silver resistance duplicates were enriched across all environments, with the strongest enrichment in Barnes-Jewish Hospital (Δ_*D*−*S*_ = 56.49%), followed by Cambridge Hospital (Δ_*D*−*S*_ = 34.57%), livestock (Δ_*D*−*S*_ = 31.40%), and WwTW (Δ_*D*−*S*_ = 9.69%) isolates. Beyond copper-silver, Barnes-Jewish Hospital isolates were enriched for tellurium duplicates (Δ_*D*−*S*_ = 26.06%), whereas Cambridge Hospital exhibited a marginal trend toward enrichment in arsenic duplicates (Δ_*D*−*S*_ = 12.67%), though this was not statistically significant. WwTW isolates were also highly enriched for arsenic duplicates (Δ_*D*−*S*_ = 40.40%). Further inspection revealed that tellurium enrichment in Barnes-Jewish Hospital was driven by a single *E. coli* isolate, whereas the arsenic duplication signal in Cambridge Hospital was limited to *Klebsiella pneumoniae* and *Citrobacter freundii*, which were not present in the Barnes-Jewish Hospital dataset. Together, these patterns indicate that duplication of ARGs and MRGs is both environment- and class-specific, reflecting unique selective pressures across sources.

## Discussion

The findings of this study show that resistance gene abundance is strongly structured by environmental selective pressures across clinical, agricultural, and wastewater settings. At this broad scale, antibiotic and metal resistance follow distinct abundance patterns shaped by the dominant stressors present in each environment, as evident from genome-level distributions of resistance gene counts and duplication frequencies. Gene duplication has historically been viewed as a general consequence of selection for increased resistance, yet this assumption has rarely been evaluated across environmental contexts. Our results demonstrate that duplication carries distinct, source-specific signatures and point to the importance of gene-level mechanisms, including association with MGEs, in maintaining and propagating resistance under local selective pressures.

Most ARGs and MRGs, irrespective of copy number, were located on plasmids, the primary vehicles for HGT of resistance genes between cells. Transposons and ISs also carried a substantial fraction of genes, consistent with their roles in facilitating gene duplication and mobilization onto plasmids or chromosomes (Partridge et al., 2018). Accordingly, the overall distribution of MGEs aligns with the well-established functions of plasmids, transposons, and ISs in the dissemination of resistance genes. Despite their well-documented role in facilitating HGT, the proportion of genes mobilized by ICEs and MITEs was negligible. Similar patterns have been reported in other studies using MobileElementFinder, in which intact ICEs and MITEs are detected at low frequencies relative to plasmids and other MGEs (Morales et al., 2023; Quijada et al., 2025). While our earlier analyses across all genes suggested that duplicate genes may have greater mobilization potential (Table S1), this pattern was observed only for ARGs in Cambridge Hospital and livestock isolates, and mobilization rates were similar between single-copy and duplicate ARGs and MRGs in other settings (Fig. 3). We hypothesize that the absence of a consistent copy-number effect reflects the high baseline mobility of ARGs and MRGs, which are frequently associated with plasmids and other MGEs (Smillie et al., 2010; Partridge et al., 2018). Thus, duplication may be a more informative predictor of mobilization for genes that lack an inherent affinity for MGEs.

Having established broader patterns of gene duplication and mobilization, we next examined enrichment within specific resistance classes. In Barnes-Jewish Hospital, duplicate genes conferring *β*-lactam resistance were enriched (Fig. 4A; Table S2), consistent with the ESBL-producing phenotype of these isolates. In contrast, Cambridge Hospital exhibited significant enrichment only in aminoglycoside–quinolone duplicate genes, rather than across multiple resistance classes. Although one might expect carbapenem resistance to coincide with enrichment of *β*-lactam duplicates, this pattern was not observed. Instead, the enrichment of aminoglycoside–quinolone-associated duplicates may reflect co-selection between carbapenem and aminoglycoside resistance (Smriti et al., 2025), driven in part by the use of combination therapies for multidrug-resistant infections (Fatsis-Kavalopoulos et al., 2022; Thy et al., 2023; Clark and Burgess, 2024). These results suggest that resistance phenotypes alone do not fully predict which resistance classes are enriched, underscoring the role of co-selection in shaping resistance profiles (Pouwels et al., 2019).

In non-clinical settings, distinct patterns of resistance class enrichment were observed. Livestock isolates showed significant enrichment of aminoglycoside ARGs (Fig. 4A; Table S2), a pattern that aligns with the widespread use of aminoglycosides in veterinary medicine and animal husbandry to treat severe bacterial infections (van Duijkeren et al., 2019; Nowacka-Kozak et al., 2023). In addition to extensive use, aminoglycosides persist in animal tissues and can accumulate at high residue levels (Kaufmann et al., 2012), imposing prolonged selective pressure that favors the maintenance and duplication of resistance genes. In WwTW isolates, the presence of duplicate ARGs is not unexpected, given the well-documented influx of antibiotics into wastewater from hospitals, pharmaceutical manufacturing, and community use, as well as the frequent detection of resistant strains in these environments (Karkman et al., 2018; Nadimpalli et al., 2020; Mutuku et al., 2022). However, the observed aminoglycoside and tetracycline duplication signals were driven by a small number of genomes, primarily *E. coli* and *K. pneumoniae* species, rather than reflecting a broad, environment-wide pattern. This suggests that while resistance itself is widespread in wastewater, duplication may depend on strain-specific properties, gene context, or recent evolutionary history. The selective forces maintaining particular duplicated resistance genes in WwTW therefore remain unclear and may reflect a combination of upstream inputs and local ecological conditions, consistent with models showing that resistance dynamics in wastewater are shaped by interacting ecological and evolutionary processes rather than uniform selection (Sutradhar et al., 2021).

Similar to ARGs, patterns of MRG duplication were closely tied to environmental sources of metal exposure. Copper–silver resistance duplicates were enriched across all four environments (Fig. 4B; Table S2). In hospital settings, this enrichment coincides with the incorporation of copper into high-touch surfaces to reduce pathogen transmission and the widespread use of silver in wound dressings and medical devices such as prosthetics, breathing tubes, and catheters (Lansdown, 2006; Grass et al., 2011). Copper–silver resistance enrichment was also observed in livestock isolates, where heavy metals are prevalent in agricultural soils due to mineral fertilizers, pesticides, and atmospheric deposition (Padhye et al., 2023; Afzal and Mahreen, 2024). Notably, both clinical and agricultural environments are subject to sustained antibiotic selective pressure, which may promote MRG persistence through the co-selection of metal and antibiotic resistance genes (Seiler and Berendonk, 2012; Pal et al., 2015). In WwTW isolates, copper–silver resistance may persist under conditions of chronic but diffuse metal exposure, which are sufficient to maintain resistance genes but are likely weaker and more variable than the sustained, high-intensity exposures characteristic of hospital and agricultural settings.

Beyond copper–silver resistance, other metal resistance classes exhibited more environment-specific patterns of duplications. Arsenic resistance duplicates were enriched in WwTW isolates (Fig. 4B; Table S2), a pattern that likely reflects the natural occurrence of arsenic in soils, minerals, and geothermal deposits (Herath et al., 2016). Furthermore, arsenic contamination in water may also derive from industrial and mining activities, as well as the historical and ongoing use of arsenic-containing compounds in agriculture and animal husbandry (Punshon et al., 2017; Zhuang et al., 2023). Because arsenic is highly toxic to most bacteria and its derivatives persist in aquatic environments, strong selective pressure favors the maintenance and amplification of arsenic detoxification and resistance genes in these settings (Rosen, 2002; Yang and Rosen, 2016). Arsenic resistance duplicates were also detected in Cambridge Hospital isolates but did not meet the threshold for statistical enrichment and were absent in Barnes-Jewish Hospital; this difference reflects species composition, as these duplicates occurred predominantly in *K. pneumoniae*, which was not represented in the Barnes dataset. Similar arsenic resistance genomic features have been documented in multidrug-resistant *K. pneumoniae* isolates from clinical case studies (Foysal et al., 2025). In contrast, enrichment of tellurium resistance duplicates in Barnes-Jewish Hospital was attributable to a single *E. coli* isolate, indicating that this signal reflects a sampling artifact rather than an environment-wide selective pattern.

Taken together, these results show that duplicate ARGs and MRGs do not arise or spread uniformly across environments but instead emerge from the combined effects of environmental exposure, gene mobility, and species composition. Clinical, agricultural, and wastewater settings each impose distinct selective pressures that shape which resistance determinants are duplicated, mobilized, and ultimately enriched. At the same time, several limitations should be considered when interpreting these patterns. Resistance gene annotation depends on curated databases that are biased toward well-characterized clinical pathogens and resistance mechanisms, potentially underrepresenting resistance determinants present in non-clinical or poorly studied taxa. In addition, the detection and classification of MGEs remain challenging, particularly for fragmented, degraded, or atypical elements, which may lead to underestimation of certain MGE classes such as ICEs or MITEs. Finally, species composition varied across environments, and some duplication signals were driven by a limited number of taxa, complicating the separation of environment-wide selective effects from lineage-specific dynamics. Despite these constraints, the consistent, environment-associated patterns observed across multiple resistance classes underscore the importance of ecological context in shaping how resistance genes persist, duplicate, and circulate within microbial communities.

## Methods

### Collection of bacterial genomes

We assembled a dataset of complete bacterial genomes from four sources representing distinct ecological contexts: (1) extended-spectrum *β*-lactamase (ESBL)-producing *E. coli* isolates from Barnes-Jewish Hospital (Mahmud et al., 2022), (2) carbapenem-producing isolates from Cambridge University Hospitals NHS Foundation Trust (Roberts et al., 2023), and (3) Enterobacterales isolates from livestock and WwTW (Shaw et al., 2021).

ESBL-producing isolates were recovered from blood and urine specimens collected from patients admitted to the medical and/or oncology wards and intensive care units of the Barnes-Jewish Hospital between June 2016 and December 2019. A total of 149 isolates from 129 patients were sequenced using Illumina short-read and Oxford Nanopore long-read technologies to generate hybrid assemblies. All assemblies are available in NCBI GenBank under BioProject accesssion PRJNA824420.

Carbapenem-producing Gram-negative bacterial isolates were collected between 2014 and 2020 at Cambridge University Hospitals from clinical diagnostic samples and screening samples (rectal swab or stool) obtained from hospital inpatients, as well as from samples referred by local general practices. *E. coli* and *K. pneumoniae* were the most common species identified. A total of 85 samples from 65 patients underwent whole-genome sequencing using Illumina short-read and Oxford Nanopore long-read platforms to produce hybrid assemblies. Only assemblies labeled as “complete” were included in this study. Genomes are available in the European Nucleotide Archive (ENA) under project accession PRJEB31034.

Livestock- and wastewater-derived Enterobacterales, primarily *E. coli*, were obtained from cattle, pig, and sheep fecal samples collected from 14 farms and from influent and effluent samples collected in 2017 from five WwTWs across the United Kingdom. Sequencing was performed using Illumina short-read and Oxford Nanopore long-read technologies to generate hybrid assemblies. A total of 495 livestock-derived and 204 wastewater-derived complete genomes were included in our analysis. Sequencing data and assemblies are available in NCBI under BioProject accession PRJNA605147.

### Identification of duplicate genes, MGEs, ARGs, and MRGs

Duplicate genes were identified using an all vs. all BLASTp search of translated coding sequences within each genome assembly. Results were filtered to retain only BLASTp hits with sequence identities greater than 85%, alignment lengths greater than 85%, bit scores greater than 50%, and e-values less than 10^−10^. Hits that did not meet all of these criteria were excluded from further analysis. When a subject sequence matched multiple queries, only the highest-scoring alignment was retained to avoid inflating duplicate counts.

MGEs were annotated using multiple complementary tools. Integrative elements were identified with MobileElementFinder v1.1.2 (Johansson et al., 2021), which incorporates data from the ISfinder, Transposon Registry, and ICEberg databases. This tool identifies insertion sequences (ISs), composite transposons, miniature inverted-repeat transposable elements (MITEs), integrative conjugative elements (ICEs), cis-mobilizable elements, and integrative mobilizable elements. Plasmid replicons were identified using PlasmidFinder v2.1.6 (Carattoli et al., 2014), and MOB-recon from MOB-suite v3.1.8 (Robertson and Nash, 2018) was used to differentiate plasmid-derived from chromosomal contigs. Prophage sequences were identified using Phigaro v2.4.0 (Starikova et al., 2020).

ARGs and MRGs were identified using AMRFinderPlus v3.12.8 (Feldgarden et al., 2021). Analyses were performed with the `--protein` option to enable protein-based searches and the `--gff` option to assign genomic coordinates to the provided protein sequences. The `--plus` option was enabled as well to include detection of virulence factors and stress response genes. ARGs were extracted from the output file by selecting entries with “core” in the “Scope” field and “AMR” in both the “Element type” and “Element subtype” fields. Similarly, MRGs were extracted by selecting entries with “Plus” in the “Scope” field, “Stress” in the “Element type” field, and “Metal” in the “Element subtype” field.

### Detection of MGE-associated duplicate genes, ARGs, and MRGs

For each genome assembly, we extracted the subject IDs of duplicate genes from the BLASTp output (see *Identification of duplicate genes, MGEs, ARGs and MRGs*). These IDs were mapped to the genomic feature format (GFF) file of the associated reference assembly to extract the start and end coordinates and contig IDs for each coding region. The same procedure was used to assign genomic features to protein sequences identified in the MobileElementFinder analysis. Because both Mob-Suite and Phigaro accept GFF files as input, genomic features for plasmids and prophages were directly available and required no additional processing.

A duplicate gene was classified as plasmid-associated if it was located on a contig labeled as “plasmid” by MOB-recon. A duplicate gene was classified as prophage-associated if it was located on the same contig and overlapped genomic coordinates with a prophage. For MGEs detected by MobileElementFinder, genes located within 31 kbp of an MGE were classified as associated and therefore potentially mobilized. This 31 kbp threshold corresponds to the length of the longest composite transposon (Tn6108) from Enterobacteriaceae in the MobileElementFinder database and was used to define the genomic window within which genes were considered to have the potential to be mobilized by surrounding integrating MGEs.

AMRFinderPlus (Feldgarden et al., 2021) reports the genomic coordinates and contig locations for each ARG and MRG. Using these outputs, we applied the same approach described above to identify associations between ARGs/MRGs and MGEs. In addition, we performed a targeted analysis restricted to duplicate ARGs and MRGs, which were defined as entries with protein IDs matching those in the list of duplicate gene protein IDs for each genome assembly.

## Statistical analyses

All statistical analyses were performed in R (R Core Team, 2025) with the RStudio IDE (Posit team, 2025).

Pairwise differences in ARG and MRG abundance across environments were evaluated using Mann–Whitney U tests (Mann and Whitney, 1947), implemented via the `wilcox.test()` function in the `stats` package (R Core Team, 2025). This nonparametric test compares ranked observations between two groups and evaluates whether their distributions differ under the null hypothesis of identical distributions. Tests were applied using the formula gene count ∼ source, where the response variable corresponded to gene counts per genome and the grouping factor represented the environment. To account for multiple pairwise comparisons, *p*-values were adjusted using the Benjamini–Hochberg procedure, and statistical significance was defined as an adjusted *p* < 0.05.

Environmental variation in gene duplication was evaluated using genome-level presence–absence data. For ARGs and MRGs, genomes were classified according to whether they contained at least one duplicate gene, and 2 × 4 contingency tables were constructed comparing duplicate presence versus absence across Barnes-Jewish Hospital, Cambridge Hospital, livestock, and WwTW isolates. Differences among environments were tested using Fisher’s exact tests (Fisher, 1922), using the `fisher.test()` function in the `stats` package (R Core Team, 2025), which assess the null hypothesis that duplicate gene presence is independent of environment. To directly compare ARG and MRG duplication within the same genomes, McNemar’s tests (McNemar, 1947) were performed using paired presence–absence data, with 2 × 2 contingency tables constructed for each environment using the `mcnemar.test()` function in the `stats` package (R Core Team, 2025). McNemar’s test evaluates the null hypothesis that the probabilities of ARG and MRG duplication are equal within genomes, accounting for the non-independent nature of these resistance determinants.

Differences in the distribution of duplicate genes between MGEs and non-mobile regions were evaluated using two-sided exact binomial tests, implemented via the `binom.test()` function in the `stats` package (R Core Team, 2025). For each sample, *x* represented the number of duplicates located within MGEs, *n* the total number of genes within MGEs, and *p* the proportion of duplicates located outside MGEs. Tests assessed whether the observed frequency of duplicates within MGEs differed from expectations based on the background proportion. Benjamini–Hochberg-adjusted *p*-values were computed using the `p.adjust(`) function (R Core Team, 2025), with significance was defined as an adjusted *p <* 0.05.

Resistance class enrichment among duplicate genes was examined by constructing 2 × 2 contingency tables for each ARG and MRG class within each environment, comparing the number of duplicate and single-copy genes assigned to a given class versus all other classes. Differences in class representation between duplicate and single-copy genes were evaluated using two-sided Fisher’s exact tests, testing the null hypothesis that resistance class membership is independent of gene copy status. For each class, the magnitude and direction of enrichment were quantified using the difference in proportions between duplicate and single-copy genes (Δ_*D*−*S*_). Classes with positive Δ_*D*−*S*_ and statistically significant Fisher’s test results were classified as enriched among duplicates (Table S2). To account for multiple testing across resistance classes, *p*-values were adjusted using the Benjamini–Hochberg procedure, and statistical significance was defined as an adjusted *p* < 0.05.

## Data availability

All bacterial genome assemblies analyzed in this study are publicly available and were obtained from previously published datasets. Complete genome assemblies from Barnes-Jewish Hospital isolates are available through NCBI under BioProject accession PRJNA824420. Genomes from Cambridge University Hospitals are available through the European Nucleotide Archive under project accession PRJEB31034. Livestock- and wastewater-derived Enterobacterales genomes are available through NCBI under BioProject accession PRJNA605147.

Processed datasets and analysis scripts are available at https://github.com/traneric/gene-duplication-and-mobility.

## Acknowledgments

This work was supported by National Institutes of Health grant R35GM142438 and National Science Foundation grant DBI-2130666.

## References

A. Afzal and N. Mahreen. Emerging insights into the impacts of heavy metals exposure on health, reproductive and productive performance of livestock. Frontiers in Pharmacology, 15:1273503, 2024.

S. Akhter, M. A. Bhat, S. Ahmed, and W. A. Siddiqui. Antibiotic residue contamination in the aquatic environment, sources and associated potential health risks. Environmental Geochemistry and Health, 46(3):1429–1454, 2024.

D. I. Andersson and D. Hughes. Gene amplification and adaptive evolution in bacteria. Annual Review of Genetics, 43:167–195, 2009.

S. Arredondo-Alonso, R. J. Willems, W. van Schaik, and A. C. Schürch. On the (im)possibility of reconstructing plasmids from whole-genome short-read sequencing data. Microbial Genomics, 3: e000128, 2017.

C. Baker-Austin, M. S. Wright, R. Stepanauskas, and J. V. McArthur. Co-selection of antibiotic and metal resistance. Trends in Microbiology, 14:176–182, 2006.

T. U. Berendonk, C. M. Manaia, C. Merlin, D. Fatta-Kassinos, E. Cytryn, F. Walsh, H. Bürgmann, H. Sørum, M. Norström, M.-N. Pons, N. Kreuzinger, P. Huovinen, S. Stefani, T. Schwartz,V. Kisand, F. Baquero, and J. L. Martinez. Tackling antibiotic resistance: the environmental framework. Nature Reviews Microbiology, 13:310–317, 2015.

M. S. Bratlie, J. Johansen, B. T. Sherman, D. W. Huang, R. A. Lempicki, and F. Drabløs. Gene duplications in prokaryotes can be associated with environmental adaptation. BMC Genomics, 11:588, 2010.

M. R. Bruins, S. Kapil, and F. W. Oehme. Microbial resistance to metals in the environment. Ecotoxicology and Environmental Safety, 45:198–207, 2000.

A. Carattoli, E. Zankari, A. García-Fernández, M. Voldby Larsen, O. Lund, L. Villa, F. Møller Aarestrup, and H. Hasman. In silico detection and typing of plasmids using plasmidfinder and plasmid multilocus sequence typing. Antimicrobial Agents and Chemotherapy, 58(7):3895–3903, 2014.

J. A. Clark and D. S. Burgess. Comparing the activity of broad-spectrum beta-lactams in combination with aminoglycosides against vim-producing enterobacteriaceae. Microbiology Spectrum, 12(1):e03876–23, 2024.

V. Economou and P. Gousia. Agriculture and food animals as a source of antimicrobial-resistant bacteria. Infection and Drug Resistance, 8:49–61, 2015.

N. Fatsis-Kavalopoulos, L. Roelofs, and D. I. Andersson. Potential risks of treating bacterial infections with a combination of -lactam and aminoglycoside antibiotics: A systematic quantification of antibiotic interactions in e. coli blood stream infection isolates. eBioMedicine, 78:103981, 2022.

M. Feldgarden, V. Brover, N. Gonzalez-Escalona, J. G. Frye, J. Haendiges, D. H. Haft, M. Hoffmann, J. B. Pettengill, A. B. Prasad, G. E. Tillman, G. H. Tyson, and W. Klimke. Amrfinderplus and the reference gene catalog facilitate examination of the genomic links among antimicrobial resistance, stress response, and virulence. Scientific Reports, 11:12728, 2021.

R. A. Fisher. On the interpretation of χ^2^ from contingency tables, and the calculation of p. Journal of the Royal Statistical Society, 85(1):87–94, 1922.

M. J. Foysal, F. Momtaz, A. M. M. A. Chowdhury, A. A. Tanni, A. Salauddin, M. Z. Hasan, S. Akter, M. A. Rahman, M. W. Rahman, D. Barua, M. M. Rahman, E. Rousham, A. Rahman, M. M. Rahman, and M. B. Hossain. Whole-genome analysis of multidrug-resistant Klebsiella pneumoniae kp04 reveals distinctive antimicrobial and arsenic-resistance genomic features: A case study from bangladesh. Current Microbiology, 82:22, 2025.

B. F. Gillieatt and N. V. Coleman. Unravelling the mechanisms of antibiotic and heavy metal resistance co-selection in environmental bacteria. FEMS Microbiology Reviews, 48:fuae017, 2024.

G. Grass, C. Rensing, and M. Solioz. Metallic copper as an antimicrobial surface. Applied and Environmental Microbiology, 77(5):1541–1547, 2011.

I. Herath, M. Vithanage, J. Bundschuh, J. P. Maity, and P. Bhattacharya. Natural arsenic in global groundwaters: Distribution and geochemical triggers for mobilization. Current Pollution Reports, 2(1):68–89, 2016.

M. H. K. Johansson, V. Bortolaia, S. Tansirichaiya, F. M. Aarestrup, A. P. Roberts, and T. N. Petersen. Detection of mobile genetic elements associated with antibiotic resistance in salmonella enterica using a newly developed web tool: Mobileelementfinder. Journal of Antimicrobial Chemotherapy, 76(1):101–109, 2021.

A. Karkman, T. T. Do, F. Walsh, and M. P. J. Virta. Antibiotic-resistance genes in waste water. Trends in Microbiology, 26(3):220–228, 2018.

A. Kaufmann, P. Butcher, and K. Maden. Determination of aminoglycoside residues by liquid chromatography and tandem mass spectrometry in a variety of matrices. Analytica Chimica Acta, 711:46–53, 2012.

C. Kingsford, M. C. Schatz, and M. Pop. Assembly complexity of prokaryotic genomes using short reads. BMC Bioinformatics, 11:21, 2010.

F. A. Kondrashov. Gene duplication as a mechanism of genomic adaptation to a changing environment. Proceedings of the Royal Society B: Biological Sciences, 279:5048–5057, 2012.

R. Kumavath, P. Gupta, E. R. Tatta, M. S. Mohan, S. A. Salim, and S. Busi. Unraveling the role of mobile genetic elements in antibiotic resistance transmission and defense strategies in bacteria. Frontiers in Systems Biology, 5:1557413, 2025.

A. B. G. Lansdown. Silver in health care: antimicrobial effects and safety in use. Current Problems in Dermatology, 33:17–34, 2006.

D. G. J. Larsson and C.-F. Flach. Antibiotic resistance in the environment. Nature Reviews Microbiology, 20:257–269, 2022.

L.-G. Li, Y. Xia, and T. Zhang. Co-occurrence of antibiotic and metal resistance genes revealed in complete genome collection. The ISME Journal, 11:651–662, 2017.

C. Llor and L. Bjerrum. Antimicrobial resistance: Risk associated with antibiotic overuse and initiatives to reduce the problem. Therapeutic Advances in Drug Safety, 5(6):229–241, 2014.

A. J. Lopatkin, H. R. Meredith, J. K. Srimani, C. Pfeiffer, R. Durrett, and L. You. Persistence and reversal of plasmid-mediated antibiotic resistance. Nature Communications, 8:1689, 2017.

R. Maddamsetti, Y. Yao, T. Wang, J. Gao, V. T. Huang, G. S. Hamrick, H.-I. Son, and L. You. Duplicated antibiotic resistance genes reveal ongoing selection and horizontal gene transfer in bacteria. Nature Communications, 15:1449, 2024.

B. Mahmud, M. A. Wallace, K. A. Reske, K. Alvarado, C. E. Muenks, D. A. Rasmussen, C.-A. D. Burnham, C. Lanzas, E. R. Dubberke, and G. Dantas. Epidemiology of plasmid lineages mediating the spread of extended-spectrum beta-lactamases among clinical Escherichia coli. mSystems, 7(3):e00519–22, 2022.

H. B. Mann and D. R. Whitney. On a test of whether one of two random variables is stochastically larger than the other. The Annals of Mathematical Statistics, 18(1):50–60, 1947.

C. Manyi-Loh, S. Mamphweli, E. Meyer, and A. Okoh. Antibiotic use in agriculture and its consequential resistance in environmental sources: Potential public health implications. Molecules, 23(4), 2018.

Q. McNemar. Note on the sampling error of the difference between correlated proportions or percentages. Psychometrika, 12(2):153–157, 1947.

G. Morales, B. Abelson, S. Reasoner, J. Miller, A. M. Earl, M. Hadjifrangiskou, and J. Schmitz. The role of mobile genetic elements in virulence factor carriage from symptomatic and asymptomatic cases of Escherichia coli bacteriuria. Microbiology Spectrum, 11(3):e04710–22, 2023.

C. Mutuku, Z. Gazdag, and S. Melegh. Occurrence of antibiotics and bacterial resistance genes in wastewater: resistance mechanisms and antimicrobial resistance control approaches. World Journal of Microbiology and Biotechnology, 38(7):152, 2022.

M. L. Nadimpalli, S. J. Marks, M. C. Montealegre, R. H. Gilman, A. Ta, L. Y. Hsu, D. Mertz, G. Trueba, A. B. Bowen, P. Rattanaumpawan, S. Shrestha, K. Phetxumphou, N. Noparatnaraporn, C. Lay, M. Cheesman, J. Jang, B. Jeon, J. E. Koehler, T. Stinear, A. Benten, and R. Laxminarayan. Urban informal settlements as hotspots of antimicrobial resistance and the need to curb environmental transmission. Nature Microbiology, 5:787–795, 2020.

E. Nowacka-Kozak, A. Gajda, and M. Gbylik-Sikorska. Analysis of aminoglycoside antibiotics: A challenge in food control. Molecules, 28(12):4595, 2023.

H. Ochman, J. G. Lawrence, and E. A. Groisman. Lateral gene transfer and the nature of bacterial innovation. Nature, 405:299–304, 2000.

S. Ohno. Evolution by Gene Duplication. Springer Berlin, Heidelberg, 1 edition, 1970.

L. P. Padhye, T. Jasemizad, S. Bolan, O. V. Tsyusko, J. M. Unrine, B. K. Biswal, R. Balasubramanian, Y. Zhang, T. Zhang, J. Zhao, Y. Li, J. Rinklebe, H. Wang, K. H. M. Siddique, and N. Bolan. Silver contamination and its toxicity and risk management in terrestrial and aquatic ecosystems. Science of The Total Environment, 871:161988, 2023.

A. Pal, S. Bhattacharjee, J. Saha, M. Sarkar, and P. Mandal. Bacterial survival strategies and responses under heavy metal stress: a comprehensive overview. Critical Reviews in Microbiology, 48:1–32, 2022.

C. Pal, J. Bengtsson-Palme, E. Kristiansson, and D. G. J. Larsson. Co-occurrence of resistance genes to antibiotics, biocides and metals reveals novel insights into their co-selection potential. BMC Genomics, 16:964, 2015.

S. R. Partridge, S. M. Kwong, N. Firth, and S. O. Jensen. Mobile genetic elements associated with antimicrobial resistance. Clinical Microbiology Reviews, 31:e00088–17, 2018.

Posit team. RStudio: Integrated Development Environment for R. Posit Software, PBC, Boston, MA, 2025.

K. B. Pouwels, B. Muller-Pebody, T. Smieszek, S. Hopkins, and J. V. Robotham. Selection and co-selection of antibiotic resistances among escherichia coli by antibiotic use in primary care: An ecological analysis. PLOS ONE, 14(10):e0218134, 2019.

T. Punshon, B. P. Jackson, A. A. Meharg, T. Warczack, K. Scheckel, and M. L. Guerinot. Understanding arsenic dynamics in agronomic systems to predict and prevent uptake by crop plants. Science of the Total Environment, 581-582:209–220, 2017. doi: 10.1016/j.scitotenv.2016.12.111.

N. M. Quijada, J. F. Cobo-Díaz, V. Valentino, C. Barcenilla, F. De Filippis, R. Cabrera-Rubio, N. Carlino, F. Pinto, M. Dzieciol, I. Calvete-Torre, C. Sabater, F. Rubino, S. Knobloch, S. Skirnisdóttir, L. Ruiz, M. López, M. Prieto, V. T. Marteinsson, A. Margolles, N. Segata, P. D. Cotter, M. Wagner, D. Ercolini, and A. Alvarez-Ordóñez. The food-associated resistome is shaped by processing and production environments. Nature Microbiology, 10(8):1854–1867, 2025.

R Core Team. R: A Language and Environment for Statistical Computing. R Foundation for Statistical Computing, Vienna, Austria, 2025.

L. W. Roberts, D. A. Enoch, F. Khokhar, G. A. Blackwell, H. Wilson, B. Warne, T. Gouliouris, Z. Iqbal, and M. E. Török. Long-read sequencing reveals genomic diversity and associated plasmid movement of carbapenemase-producing bacteria in a uk hospital over 6 years. Microbial Genomics, 9(7):mgen001048, 2023.

J. Robertson and J. H. E. Nash. Mob-suite: software tools for clustering, reconstruction and typing of plasmids from draft assemblies. Microbial Genomics, 4(8):e000206, 2018.

E. P. Rocha, M. Haudiquet, J. de Sousa, and M. Touchon. Selfish, promiscuous and sometimes useful: how mobile genetic elements drive horizontal gene transfer in microbial populations. Philosophical Transactions of the Royal Society B: Biological Sciences, 377(1843):20210234, 2022.

B. P. Rosen. Biochemistry of arsenic detoxification. FEBS Letters, 529(1):86–92, 2002. doi: 10.1016/S0014-5793(02)03186-1.

A. San Millan and R. C. MacLean. Fitness costs of plasmids: a limit to plasmid transmission. Microbiology Spectrum, 5(5):MTBP0011–2016, 2017.

L. Sandegren and D. I. Andersson. Bacterial gene amplification: implications for the evolution of antibiotic resistance. Nature Reviews Microbiology, 7:578–588, 2009.

C. Seiler and T. U. Berendonk. Heavy metal driven co-selection of antibiotic resistance in soil and water bodies impacted by agriculture and aquaculture. Frontiers in Microbiology, 3:399, 2012.

L. P. Shaw, K. K. Chau, J. Kavanagh, M. AbuOun, E. Stubberfield, H. S. Gweon, L. Barker, G. Rodger, M. J. Bowes, R. Hubbard, T. E. A. Peto, D. W. Crook, D. S. Read, M. F. Anjum, A. S. Walker, and N. Stoesser. Niche and local geography shape the pangenome of wastewater- and livestock-associated Enterobacteriaceae. Science Advances, 7(7):eabe3868, 2021.

C. Smillie, M. P. Garcillán-Barcia, M. V. Francia, E. P. C. Rocha, and F. de la Cruz. Mobility of plasmids. Microbiology and Molecular Biology Reviews, 74(3):434–452, 2010.

S. Smriti, G. Verma, S. Pradhan, N. Singh, S. S. Panda, I. Mohapatra, D. Pattnaik, R. K. Dash, and L. Das. Co-occurrence of genes encoding carbapenem resistance and aminoglycoside resistance in clinical isolates of enterobacterales. Drug Target Insights, 19:1–10, 2025.

E. V. Starikova, P. O. Tikhonova, N. A. Prianichnikov, C. M. Rands, E. M. Zdobnov, E. N. Ilina, and V. M. Govorun. Phigaro: high-throughput prophage sequence annotation. Bioinformatics, 36(12):3882–3884, 2020.

I. Sutradhar, C. Ching, D. Desai, M. Suprenant, E. Briars, Z. Heins, A. S. Khalil, and M. H. Zaman. Computational model to quantify the growth of antibiotic-resistant bacteria in wastewater. mSystems, 6(3):e00360–21, 2021. doi: 10.1128/mSystems.00360-21.

M. Thy, J.-F. Timsit, and E. de Montmollin. Aminoglycosides for the treatment of severe infection due to resistant gram-negative pathogens. Antibiotics, 12(5):860, 2023.

M. Tokuda and M. Shintani. Microbial evolution through horizontal gene transfer by mobile genetic elements. Microbial Biotechnology, 17:e14408, 2024.

E. van Duijkeren, C. Schwarz, D. Bouchard, B. Catry, C. Pomba, K. E. Baptiste, M. A. Moreno, M. Rantala, M. Ružauskas, P. Sanders, C. Teale, A. L. Wester, K. Ignate, Z. Kunsagi, and H. Jukes. The use of aminoglycosides in animals within the eu: development of resistance in animals and possible impact on human and animal health: a review. Journal of Antimicrobial Chemotherapy, 74(9):2480–2496, 2019.

World Health Organization. No time to wait: Securing the future from drug-resistant infections. Technical report, World Health Organization, 2019. Report to the Secretary-General of the United Nations.

H.-C. Yang and B. P. Rosen. New mechanisms of bacterial arsenic resistance. Biomedicine Journal, 39(2):59–68, 2016.

W. Zhao, W. Zeng, B. Pang, M. Luo, Y. Peng, J. Xu, B. Kan, Z. Li, and X. Lu. Oxford nanopore long-read sequencing enables the generation of complete bacterial and plasmid genomes without short-read sequencing. Frontiers in Microbiology, 14:1179966, 2023.

F. Zhuang, J. Huang, H. Li, X. Peng, L. Xia, L. Zhou, T. Zhang, Z. Liu, Q. He, F. Luo, H. Yin, and D. Meng. Biogeochemical behavior and pollution control of arsenic in mining areas: A review. Frontiers in Microbiology, 14:1043024, 2023. doi: 10.3389/fmicb.2023.1043024.

