## Supplementary Figures for "Ecological context structures duplication and mobilization of antibiotic and metal resistance genes in bacteria"

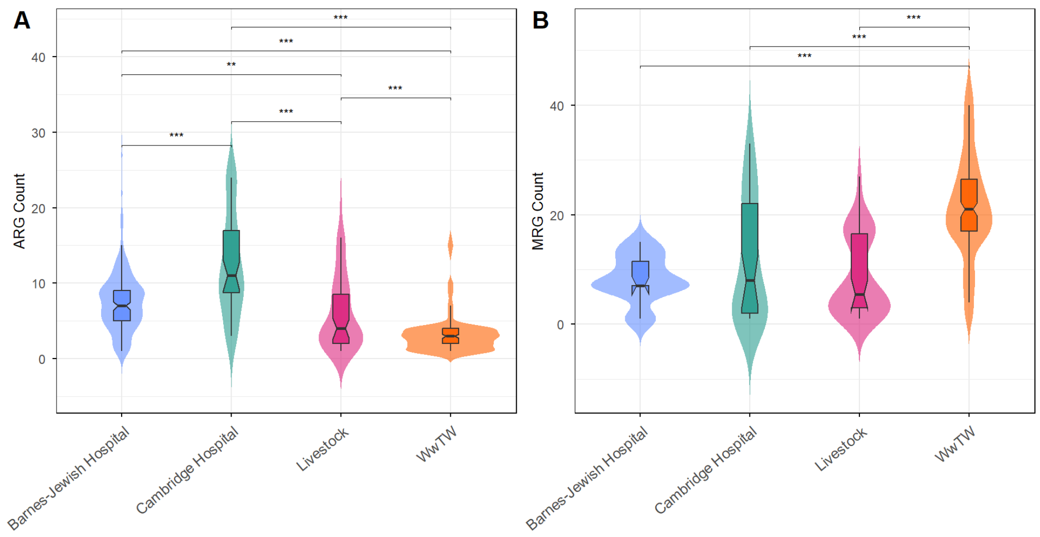


Figure S1: Abundance of ARGs (**A**) and MRGs (**B**) per genome across the four environments. Violin plots show distributions of gene counts, with embedded boxplots indicating the median and interquartile range. Analyses were performed with *E. coli* filtered from all environments except Barnes–Jewish Hospital, which consists exclusively of *E. coli* isolates and was therefore retained. $*p < 0.05$, $**p < 0.01$, $***p < 0.001$ (see *Methods*).

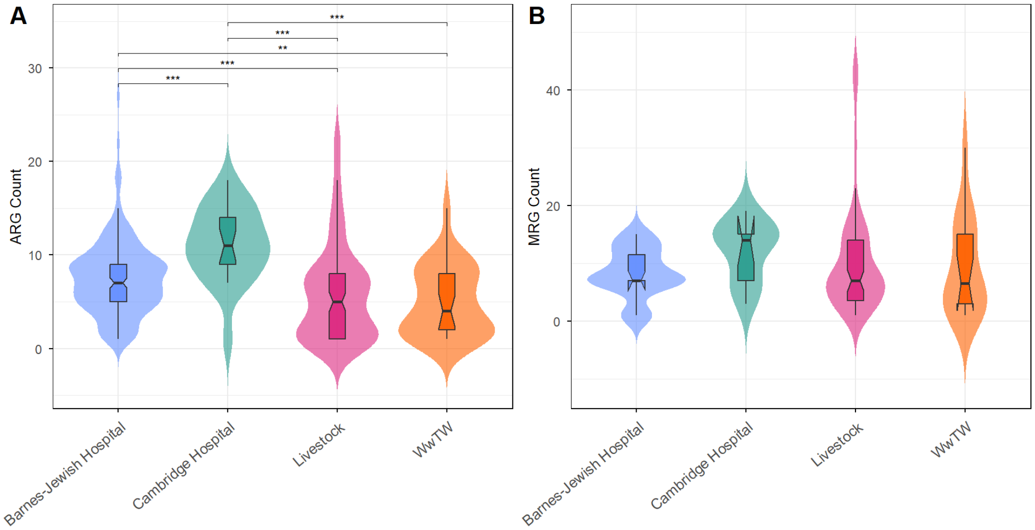


Figure S2: Abundance of ARGs (**A**) and MRGs (**B**) per genome across the four environments. Violin plots show distributions of gene counts, with embedded boxplots indicating the median and interquartile range. Analyses were performed with *E. coli*-only. $*p < 0.05$, $**p < 0.01$, $***p < 0.001$ (see *Methods*).
